# Genomic insights reveal community structure and phylogenetic associations of endohyphal bacteria and viruses in fungal endophytes

**DOI:** 10.1101/2025.02.07.637084

**Authors:** E. Escudero-Leyva, M. Belle, A. Dadkhahtehrani, J.N. Culver, M. Araya-Salas, J.P. Kutza, N. Goldson, M. Chavarría, P. Chaverri

## Abstract

**Background:** Endohyphal microbiomes, comprising bacteria and viruses, significantly influence fungal phenotypes, host fitness, and ecological interactions. Endohyphal bacterial symbionts are known to affect fungal pathogenicity, secondary metabolite production, and adaptability, yet many aspects of their diversity and interactions remain uncertain. In contrast, endohyphal viruses, despite their widespread presence, are poorly understood in terms of their diversity, ecological roles, and evolutionary relationships with fungal hosts. These gaps highlight the need for integrative studies to explore the composition, diversity, host associations, and functional roles of both bacterial and viral communities in fungi. This study aimed to (1) characterize the diversity of endohyphal bacterial and viral communities within selected fungal endophytes using genomic and transcriptomic approaches, and (2) test for host specialization through phylogenetic signals and core microbial taxa.

**Results:** Endohyphal microbial communities from 19 fungal isolates from eight fungal orders (*Amphisphaeriales, Botryosphaeriales, Diaporthales, Glomerellales, Mucorales, Pleosporales, Sordariales*, and *Xylariales*) obtained from *Fagus grandifolia* leaves were characterized. Bacterial communities exhibited high diversity and significant phylogenetic signals, with core taxa such as *Bacillales, Burkholderiales, Enterobacterales, Hyphomicrobiales*, and *Pseudomonadales*, shared across all fungal samples. Specific bacterial taxa displayed potential host specialization, including *Moraxellales, Sphingomonadales*, and *Streptosporangiaceae* for *Amphisphaeriales* fungal samples; *Enterobacterales* (e.g., *Enterobacteraceae*), *Hyphomicrobiales* (e.g., *Rhizobiaceae*), and *Micrococcales* for *Glomerellales*; and *Cytophagales* for *Diaporthales*. Viral communities were less diverse, with *Bamfordvirae* and *Heunggongvirae* (double-stranded DNA viruses) identified as core taxa in metagenomic data, while metatranscriptomic data revealed no core taxa. Surprisingly, only a few reads of double-stranded RNA mycoviruses were detected.

**Conclusions:** The findings suggest a level of host specialization in bacterial communities and a more complex, limited association for viral communities, with dsDNA dominating the endohyphal virome. This study enhances our understanding of fungal-microbe interactions and highlights the ecological and evolutionary dynamics of endohyphal microbiota. Future efforts to expand reference databases and investigate the functional roles of these microbial taxa will further elucidate their contributions to fungal biology, impacts on their plant hosts, and ecosystem processes.

## BACKGROUND

Symbiotic relationships within an ecological network can be classified as direct or indirect [1,2]. A direct relationship involves the immediate effect of one organism on another when not mediated or transmitted through a third individual. An example of a direct interaction is fungal endophytism, where fungi live within healthy plant tissues without causing disease symptoms.

Some endophytes may eventually become pathogens [3], while others remain commensals or mutualists, aiding plant growth and offering protection against diseases or abiotic stressors [4]. In contrast, an indirect relationship can be exemplified by the endohyphal microbiota— microorganisms residing within the hyphae of endophytic fungi. These microorganisms can have positive or negative effects on their fungal host, which in turn, indirectly influences plant biology [5–7]. Disentangling the plant’s holobiont and its symbiotic relationships is crucial for understanding how the presence or absence of one or multiple symbionts can trigger cascading effects, ultimately influencing the fate of plants and, hence, plant communities and ecosystems [8,9].

Despite advancements in tools to more comprehensively characterize microbial communities (e.g., metagenomics, metatranscriptomics, and bioinformatics), the full spectrum of the multi-species and multi-level symbiotic relationships within plant-associated fungi (e.g., endophytes) remains poorly known. A few studies have revealed the complexity of these endohyphal communities, which include bacteria, viruses, and, occasionally, microalgae [6,10,11]. However, basic knowledge, such as alpha or beta diversity within fungal hyphae and across fungal taxa, especially for endohyphal viruses, is limited.

Endohyphal bacteria have been studied more extensively than viruses (e.g., [12–16]). Many studies have focused on the effects they can induce in their fungal hosts [5,17]. For instance, they can reduce or increase pathogenicity [18,19], increase spore and mycotoxin production [20], heighten respiration and hyphal density [21], protect against predatory nematodes and amoebae [22], and support endophytic establishment [23,24], among others. Other studies have focused on characterizing their diversity. For example, a large-scale 16S ribosomal DNA metabarcoding study that screened hundreds of fungal isolates across several taxonomic levels revealed bacterial associations in nearly 90% of the isolates [11], in contrast to the ∼20% found by Hoffman & Arnold [13]. However, most published studies have concentrated on a few basal fungal groups (e.g., *Glomeromycota, Morteriellomycota*, and *Mucoromycota*), with even less attention given to other highly diverse fungal phyla (e.g., *Ascomycota* and *Basidiomycota*) [6], highlighting the opportunity to explore and uncover a vast array of uncharacterized diversity.

Bacteria can associate with fungi both externally and internally, via horizontal or vertical transmission, with the nature of the interaction influenced by the fungal morphology as well as the surface molecules and secreted factors of both the fungi and bacteria [25]. Therefore, one would expect that host-specificity, limited host ranges, or coevolution would be more widespread in fungi-bacteria interactions, fostering increased species diversity by promoting niche specialization and enabling resource partitioning, among other factors [26–28]. However, studies have reported that while certain bacterial groups, such as *Pseudomonadota* (*Proteobacteria*), *Actinomycetota* (*Actinobacteria*), and *Bacillota*, tend to associate with most fungi, host specificity is not always evident [11,13,14]. Some well-known examples of obligate symbiosis include *Betaproteobacteria* with *Mucoromycota* (e.g., fungus *Rhizopus* - bacterium *Mycetohabitans*), *Glomeromycota* (e.g., *Gigaspora* - ‘*Candidatus Glomeribacter gigasporum’*), and *Mortierellomycota* (e.g., *Mortierella* - *Mycoavidus*) [25].

The diversity and function of endohyphal viruses remain largely unexplored compared to endohyphal bacteria. Although most research has focused on mycoviruses in pathogenic *Ascomycota* and *Basidiomycota*, these viruses are present across all fungal lineages [29]. Recent advancements in high-throughput sequencing technologies have revealed that the fungal virome encompasses a variety of genome types, altering the previous misconception that mycoviruses were primarily double-stranded RNA (dsRNA) [30–33]. Mycoviral diversity includes more than 25 families, many of which have RNA genomes (e.g., dsRNA), and others with DNA genomes. The functions of most mycoviruses are poorly known, except for some dsRNA viruses. For example, several dsRNA viruses can confer hypovirulence, making them potential candidates for the biocontrol of mycotoxigenic or plant-pathogenic fungi [33–38]. These dsRNA hypovirulence mycoviruses can also transform non-pathogenic endophytic fungi (e.g., *Colletotrichum*, *Pestalotiopsis*, and *Sclerotinia*, among others) into highly phytopathogenic fungi [38–42].

Conversely, hypervirulence has emerged as an attractive strategy to enhance the biocontrol effectiveness of entomopathogenic fungi, such as *Beauveria bassiana* and its associated virus, ‘*Beauveria bassiana* victorivirus 1 (BbVV-1)’ [36,43].

Mycoviruses may be transmitted through fungal spores (vertical transmission) or hyphal anastomosis (horizontal transmission) [10,44], with no known extracellular entry route [45]. This lack of an extracellular entry route has been a significant challenge for the intended propagation of mycoviruses and their effective use in biological control [46]. Additionally, some viruses (e.g., dsDNA viruses) have incorporated viral elements into fungal genomes through mechanisms that are not yet fully understood, occurring either in recent or ancient (millions of years) events [47–49]. The above-mentioned transmission mechanisms suggest the potential for niche specialization and/or coevolution, which could then result in some level of host-specificity [26–28]. However, no host-specificity or preference has been reported to date.

Based on what is known of bacterial and viral transmission in fungi, we hypothesize that there is a phylogenetic association (i.e., phylogenetic signal) between the fungal host and the diversity of endohyphal bacteria and viruses. Therefore, the aims of this study were two-fold. First, we aimed to characterize the bacterial and viral genomic and transcriptomic diversity residing in the hyphae of selected endophytes isolated from *Fagus grandifolia* (American beech) using *de novo* metagenomics and metatranscriptomics. Second, with the data obtained from the first objective, we tested for phylogenetic signals and core microbial taxa. The results of this study will enhance our understanding of microbial diversity, particularly how it may influence the phenotype and genotype of the host fungus and its indirect interactions with plants. This knowledge is crucial for elucidating the complex multi-species and multi-level symbiotic relationships within plant-associated fungi and their potential applications in pathogen-specific biocontrol and ecosystem health.

## METHODS

### Fungal isolation and identification

Endophytic fungi were obtained following previously used protocols [50,51]. These were isolated from two *Fagus grandifolia* trees located in the forest adjacent to Bowie State University (39° 01’ 17.69" N; -76° 45’ 24.62" W); five healthy leaves per tree were collected. Five discs (5 mm in diameter) per leaf were excised and surface-sterilized by sequential immersion in bleach (2% for 30 seconds), ethanol (70% for 1 minute), and thoroughly rinsed with sterile distilled water. The sterilized discs were placed in Petri dishes (10 cm) containing potato dextrose agar (PDA, 15 mL, Difco, Detroit, MI, USA). Plates were incubated (25 °C), and subculturing was performed once hyphal growth began, transferring the mycelium to fresh PDA plates to establish axenic cultures.

DNA extraction was done after approximately 7 days of growth in PDA. The mycelium was harvested and processed using the PowerPlant Mini kit (Qiagen Inc., Hilden, Germany) following the manufacturer’s instructions. The internal transcribed spacers region (ITS1, 5.8S, ITS2) of the nuclear ribosomal DNA (ITS nrDNA) was amplified using primers ITS4 and ITS5 [52]. Additional gene regions were amplified to refine the identification for selected isolates: translation-elongation factor 1-α (TEF) using primers EF728f and EF2r [53], and β-tubulin (TUB) using primers Bt1 and Bt2 [54]. The PCR reaction mixture consisted of GoTaq Green Master Mix (12.5 µL; Promega Corporation, Madison, WI, USA), forward and reverse primers (1 µL of each), bovine serum albumin (BSA, 0.5 µL, ThermoFisher Scientific, Waltham, MA, USA), dimethyl sulfoxide (DMSO, 1 µL, Merck, Darmstadt, Germany), UltraPure nuclease-free water (7 µL, ThermoFisher Scientific) and the template DNA (2 µL) [55]. PCR products were purified and sequenced by Psomagen (Maryland, USA). The sequences were trimmed and aligned using BioEdit v.7.7.1 [56], and BLASTn searches were conducted against the NCBI GenBank database for each gene region. Fungal identification was based on sequences with >98% identity and >85% coverage [57], prioritizing the most complete taxonomic classification available. Newly generated sequences were deposited in GenBank (Supplementary Table S1).

Once all collected endophyte isolates were identified, we selected samples that represented the diversity found in *F. grandifolia* leaves, ensuring at least four representatives of some fungal orders and multiple isolates from the same fungal family to test for host association. After this selection, 19 isolates remained for the subsequent analyses: *Amphisphaeriales* (4 isolates), *Diaporthales* (5), *Glomerellales* (4), which were those with more representatives; and then *Botryosphaeriales* (1), *Mucorales* (1), *Pleosporales* (1), *Sordariales* (1), and *Xylariales* (2), which had fewer isolates (Supplementary Table S1). Of those, the best-represented families were *Diaporthaceae, Glomerellaceae*, and *Pestalotiopsidaceae*; *Diaporthe*, *Colletotrichum,* and *Pestalotiopsis* as their corresponding genera.

### Fungal phylogenetic analysis

To be able to test for phylogenetic signals, we first had to reconstruct a phylogeny for the fungal isolates selected. For the selected 19 samples, sequences from each gene were independently aligned using MUSCLE within MEGA v.11 [58,59]. To ensure comparable sequence lengths, the alignments were trimmed: ITS nrDNA alignment to 676 bp, TEF to 398 bp, and TUB to 404 bp. Then, the trimmed alignments were concatenated using MEGA and exported to Phylip format for phylogenetic tree reconstruction. Prior to building the tree, ModelFinder [60] was used inside IQ-TREE v.2.2.0 [61] to identify the most suitable substitution model for the concatenated dataset. To assess the robustness of the inferred relationships, 1000 bootstrap replicates were performed. This analysis was done using the Kabré HPC cluster (CeNAT-CONARE, Costa Rica). Finally, the consensus tree was visualized using FigTree v.1.4.4 (http://tree.bio.ed.ac.uk/software/figtree/) setting sample M22 (*Umbelopsis* aff. *isabellina, Mucoromycota*) as the outgroup.

### Total DNA and RNA extraction and sequencing

Following purification, fungal isolates were grown in liquid culture. Erlenmeyer flasks (250 mL) containing malt extract broth (MEB, 50 mL, Difco, Detroit, MI, USA) were inoculated with five mycelial plugs (5 mm diameter) and placed in an I24 New Brunswick Scientific incubator shaker (New Brunswick Scientific, New Jersey, USA) (25 °C, 110 rpm) for seven days. Before harvesting the mycelium, it was confirmed that the broth was clear and free of bacterial contaminants. Fresh mycelium was harvested using a Buchner funnel system, sterile filter paper, and vacuum, then dried with sterile paper towels, immediately flash-frozen with liquid nitrogen, and stored at -80 °C for a maximum of five days. Approximately 100 mg of frozen mycelia were then ground using a mortar and pestle with liquid nitrogen throughout the process. Each sample was ground separately at different times to avoid cross-contamination. Then, total DNA and RNA were extracted from the ground mycelia using commercially available kits (DNeasy Plant Pro Kit and RNeasy Plant Mini Kit; Qiagen Inc., Hilden, Germany) following the manufacturer’s instructions. Total RNA and DNA were further cleaned and concentrated using DNA and RNA Clean and Concentration kits (Zymo Research Corporation, Irvine, California, USA) according to the manufacturer’s protocol. The purified DNA and RNA samples were then stored (-80 °C). Total DNA and RNA were sent to Novogene Inc. (California, USA) for shotgun metagenomic and metatranscriptomic sequencing (NovaSeq PE150; 6Gb per sample; ribosomal depletion for RNA samples). All raw data (.fastq files) have been deposited in GenBank under BioProject PRJNA1221291.

### Bioinformatic analyses of metagenomic and metatranscriptomic data

Computational analyses were performed on the Kabré HPC Cluster. Data quality control and filtering were done by Novogene, including adapter and low-quality sequence removal, resulting in raw data quality >92% Q30. Reads were further filtered and trimmed with Seqtk v.1.4 (https://github.com/lh3/seqtk/) and BBDuk v.38.84 (https://sourceforge.net/projects/bbmap/).

Assembly of the metagenomic and metatranscriptomic data was done using metaSPAdes [62] and rnaSPAdes [63], respectively. After assembly and before running the taxonomy classifier on metagenome and metatranscriptome assemblies, contigs/transcripts were filtered to a minimum length of 500 bp. Taxonomic assignments were made with Kaiju [64], selecting the “nr” v.2023-05-10 and “virus” v.2023-05-10 databases for prokaryotes and viruses, respectively. Kaiju is considered a sensitive taxonomy classifier that uses NCBI RefSeq database and protein-based classifiers [64–66]. Kaiju was run for prokaryotic and viral contigs with metagenomic data, and only for viral transcripts for metatranscriptomic data. The latter was done to determine if this approach would better detect dsRNA viruses. Counts and taxonomy tables generated by Kaiju were manually inspected and curated to remove potential artifacts and ensure data quality.

### Transmission electron microscopy

Transmission electron microscopy (TEM) was performed with selected fungal isolates as another tool to investigate the presence of potential endohyphal bacteria and viruses. Isolates K21, M5t, and M67 were chosen for their apparent high number of anticipated viral contigs and taxa when compared to other isolates. These three isolates were cultured in MEB and shaken (∼110 rpm, 20 °C) for 7 days in the dark. A conventional fixation and embedding method was used as described previously [67] with the following specifications. Hyphae from each culture were excised in fixative (4% paraformaldehyde and 3% glutaraldehyde in sodium phosphate buffer 0.1mol/L, pH 7.1), washed three times in fixative, and stored (4 °C) for two days in the same fixative. Post-fixation was performed (1% osmium tetroxide for 2 hours) and then washed with distilled water (10–15 minutes). Dehydration of samples was performed by a series of 15-minute ethanol treatments at concentrations of 25%, 50%, 75%, and 90%, and then in absolute ethanol (1 hour). Dehydrated samples were infiltrated with absolute ethanol:Spurr resin [68] in mixtures (3:1, 1:1, 1:3 for 1 hour each), then in pure resin (12 hours). The polymerization of the resin was achieved at 60 °C for 24 hours. Fully polymerized samples were ultrathin sectioned (60–100 nm) and stained with aqueous uranyl acetate (1–2%) and lead citrate (3%) [69]. Thin sections were viewed in an HT7700 Transmission Electron Microscope (Hitachi, Japan; acceleration voltage up to 80 kV).

### Community analyses

To evaluate the composition of the endohyphal microbiota, metagenomic and metatranscriptomic data were analyzed using R v.4.3.3 [70]. First, to assess the extent of the sampling effort, taxa accumulation curves were generated with the iNEXT package using sequence abundance [71].

For beta diversity, two analyses were done: (1) with all the samples and (2) with samples with major representation to create a balanced dataset. This second dataset included isolates from *Amphisphaeriales* (K2, K5, K10, K18), *Diaporthales* (M1b, M13t, M20, M24, M32) and *Glomerelalles* (M19, M27, M48, M66). For both datasets, the count data were transformed using the Hellinger method with the “decostand” function from the vegan package v.2.6.4 [72]. To evaluate differences among fungal orders and families, as well as differences across isolates, an ANOVA was performed for the full dataset and PERMANOVA for the balanced dataset, and a multivariate test was done for both using “adonis” and “betadisper” functions from vegan.

Subsequently, non-metric multidimensional scaling (NMDS) was performed using a Bray-Curtis transformed matrix with vegan using “metaMDS” function. Additionally, indicator species analysis was performed using indicspecies package v.1.7.15 through the function “multipat” (multilevel pattern analysis) to retrieve possible bacterial or viral taxa important to the fungal orders using the species occurrence variable [73]. Then, to capture the overall structure of the community [74], a multinomial species classification was run with the function “clamtest” from the vegan package. Finally, to determine the core microbiome, taxa that were found in >80% of the samples and with a relative abundance greater than 0.1% were considered “core,” and those present in 50–79% of the samples were considered “resident” [75–78]. Relative abundances (%) were calculated and plotted using the R package microeco [79].

### Phylogenetic signal analyses

To assess a potential phylogenetic signal between endohyphal communities and their fungal hosts, a Mantel test was conducted for the full and balanced datasets in R v.4.3.3 using the vegan package [70,72]. The maximum likelihood matrix derived from the concatenated phylogenetic tree generated with IQ-TREE, alongside abundance tables for each community (bacteria and viruses) and technique (metagenomic and metatranscriptomic). The abundance tables were converted to presence/absence matrices, followed by a Jaccard transformation using the “vegdist” function from vegan. The Mantel test was then run with 10,000 permutations. To validate the model, simulations were performed using the MASS package [80] randomly assigning different combinations of endohyphal communities and fungal hosts across 1,000 replicates. This approach was used to evaluate Type I and Type II errors. The script for these analyses can be consulted in the following link (https://rpubs.com/marcelo-araya-salas/1218708).

Additionally, the Procrustean superimposition approach (Procrustes) [81] was performed using the NMDS values and the distance-based redundancy analysis (dbRDA) to evaluate the association of the communities within the order taxonomical level of fungi. This was done first by calculating the dbRDA values with the function “dbRDA” from the vegan package and then using the function “protest” in vegan.

## RESULTS

### Fungal isolate identification and phylogeny

From the leaves of two *Fagus grandifolia* trees, 97 endophytic fungal isolates were recovered: 55 *Sordariomycetes* (*Apiospora, Beltrania, Colletotrichum, Diaporthe, Neopestalotiopsis, Nigrospora, Pestalotiopsis, Tubakia*, and unidentified *Xylariales*), 41 *Dothideomycetes* (*Cladosporium, Didymosphaeria*, and *Lasiodiplodia*), and one *Mucoromycetes* (*Umbelopsis*). (results not shown). Out of those, 19 isolates (already described in the Methods) were selected; most of them *Sordariomycetes*: 4 *Amphisphaeriales*, 5 *Diaporthales*, 4 *Glomerellales*, 1 *Botryosphaeriales*, 1 *Mucorales*, 1 *Pleosporales*, 1 *Sordariales*, and 2 *Xylariales* (Supplementary Table S1). The concatenated phylogenetic tree formed mostly well-supported clades that, in general, corresponded to the fungal families and orders (Supplementary Figure S1).

### Bacterial metagenomic data analyses

#### Alpha diversity

After assembly and taxonomic classification, 7,747 metagenomic contigs were matched to bacteria using the Kaiju taxonomy classifier (Supplementary Table S2). Variability in contig counts was observed across samples, with most samples generating approximately 2,000 contigs and representing nearly 400 taxa. An exception was sample M22, which had significantly fewer contigs (Supplementary Figure S2A). When classified by fungal order, *Xylariales* was particularly notable, representing over 500 bacterial taxa across 2,000 contigs, despite being derived from only two samples. Most other orders, except *Botryosphaeriales* and *Mucorales*, comprised fewer than 300 taxa. *Botryosphaeriales* and *Mucorales* accounted for fewer than 250 taxa across 1,000 contigs (Supplementary Figure S2B). At the family level, most groups contained approximately 400 taxa, with *Umbelopsidaceae* showing the lowest number (<200). Extrapolated data suggested that *Apiosporaceae* might contain nearly 3,000 contigs, potentially representing up to 600 taxa. For other families, the data followed a similar trend, with 2,000 contigs yielding around 300 taxa (Supplementary Figure 2C). However, none of the isolates across the taxonomical classifications reached an asymptote, suggesting that bacterial richness might be even greater than observed. TEM analysis identified multiple bacteria-like cells within M5t and K21 isolates but not within M67 (Fig. 1).

**Figure 1.**
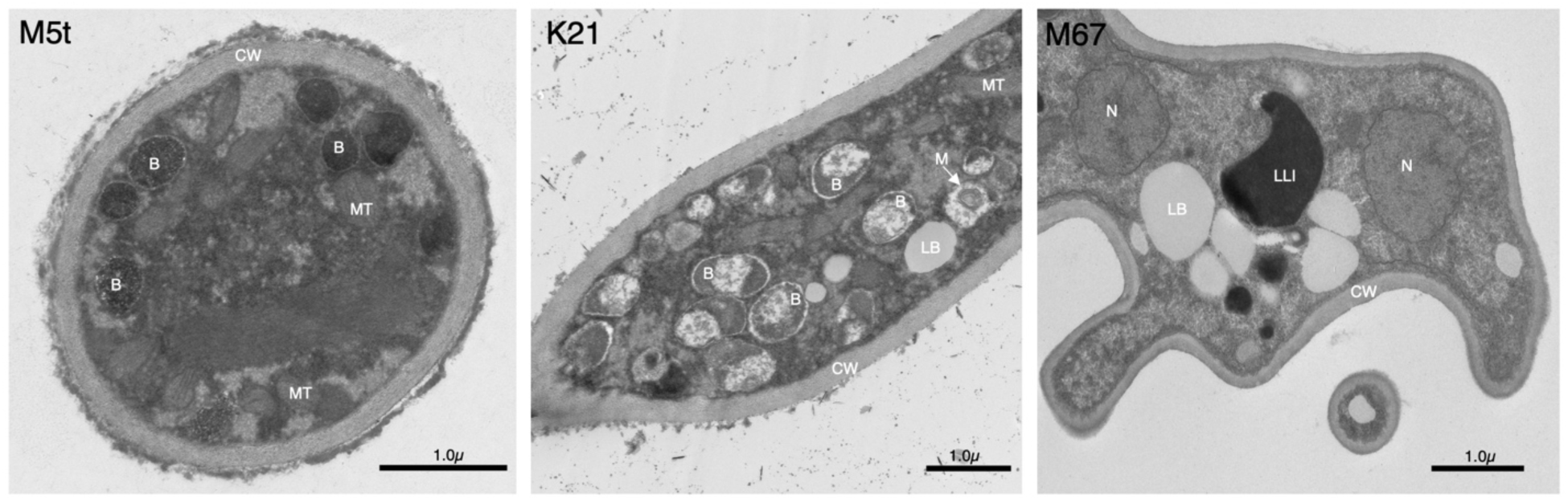
Transmission Electron Microscopy (TEM) image of hyphal cross-sections of isolates M5t, K21 and M67. *B:* Endohyphal bacteria; *CW*: Fungal cell wall; *LB*: Fungal lipidic body; LLI: Fungal Lipid-like inclusion; *M:* Bacterial mesosome; *MT*: Fungal mitochondria.

#### Bacterial community structure, composition, and phylogenetic signal across fungal hosts (metagenomics)

The analysis for the full dataset indicated a difference among the communities across the fungal orders (*F_7,11_* = 1.13, *P* = 8.013E-05) with heterogeneity across the groups (*F_7,11_* = 1.13, *P* = 0.0238). In the analysis by fungal families, the difference across the groups was maintained (*F_9,9_* = 1.80, *P* = 0.0044) but dispersion among the samples was similar (*F_9,9_* =1.80, *P* = 0.0907). The NMDS stress value (0.0838) indicated a good fit for the model and visualization by order and family correlates with the values obtained for the dispersion tests (Supplementary Figures S3A and S3B). For the balanced analyses, restricted to samples belonging to the *Amphisphaeriales*, *Diaporthales*, and *Glomerellales*, either by order or family, differences were not found (*F_2,9_* = 1.06, *P* = 0.312; *F_2,9_* = 1.15, *P* = 0.1166 respectively). Samples across these two taxonomic levels were also homogeneous (*F_2,9_* = 0.2678, *P* = 0.8776 for the order and *F_2,9_* = 0.389 *P* = 0.7532 for the family). The NMDS resulted in a good fit (stress value = 0.0829); however, consolidated and close clusters were not clearly seen by order or family in the visualization (Supplementary Figures S4A and S4B).

The bacterial community composition revealed that 14 orders, 14 families, and 10 genera were consistently present in 80–100% of the fungal samples (i.e., core); and 13 orders, 20 families, and 10 genera were present in 50–79% of the samples (i.e., resident) (Figure 2, Supplementary Tables S3–S5). Remarkably, ∼48%, 51%, and 70%, of the detected bacteria at the order, family, and genus levels, respectively, were unidentified/unclassified. The most abundant core bacterial orders across all samples were *Bacillales, Burkholderiales, Cytophagales, Enterobacterales, Hyphomicrobiales, Lactobacillales, Moraxellales, Neisseriales, Pseudomonadales*, and *Rhodobacterales*, with relative abundances ∼1.5–7% (Figures 2 and 3, Supplementary Tables S2–S5). The top five most abundant core bacterial families and genera were *Alcaligenaceae, Moraxellaceae, Rhizobiaceae, Sphingobacteriaceae*, and *Streptomycetaceae*; and *Coraliihabitans, Erythrobacter/Porphyrobacter* complex, *Pseudomonas*, *Rhizobium/Agrobacterium* complex, and *Salmonella,* respectively (Figures 2 and 3, Supplementary Tables S2–S5, Supplementary Figure S5). The top five most abundant resident bacterial orders, families, and genera across all fungal samples were *Flavobacteriales, Mycobacteriales, Propionibacteriales, Rickettsiales*, and *Sphingomonadales; Beijerinckiaceae, Lactobacillaceae, Microbacteriaceae, Morganellaceae*, and *Streptococcaceae*; and *Azorhizobium, Enterobacter, Francisella*, unidentified *Rickettsiales*, and *Vibrio*, respectively (Figures 2 and 3, Supplementary Tables S2–S5).

**Figure 2.**
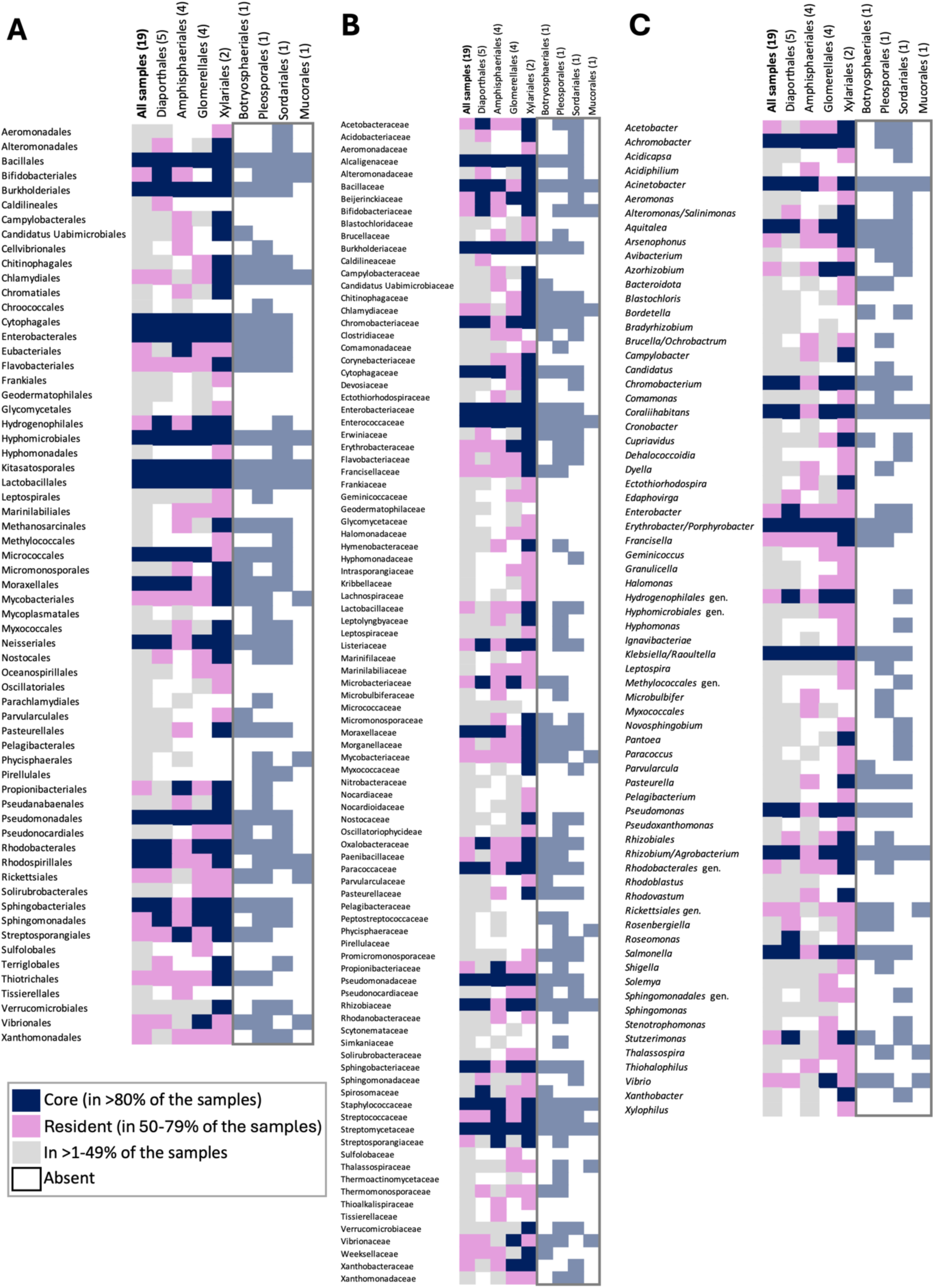
Core and resident bacterial orders (A), families (B), and genera (C) by fungal order.

**Figure 3.**
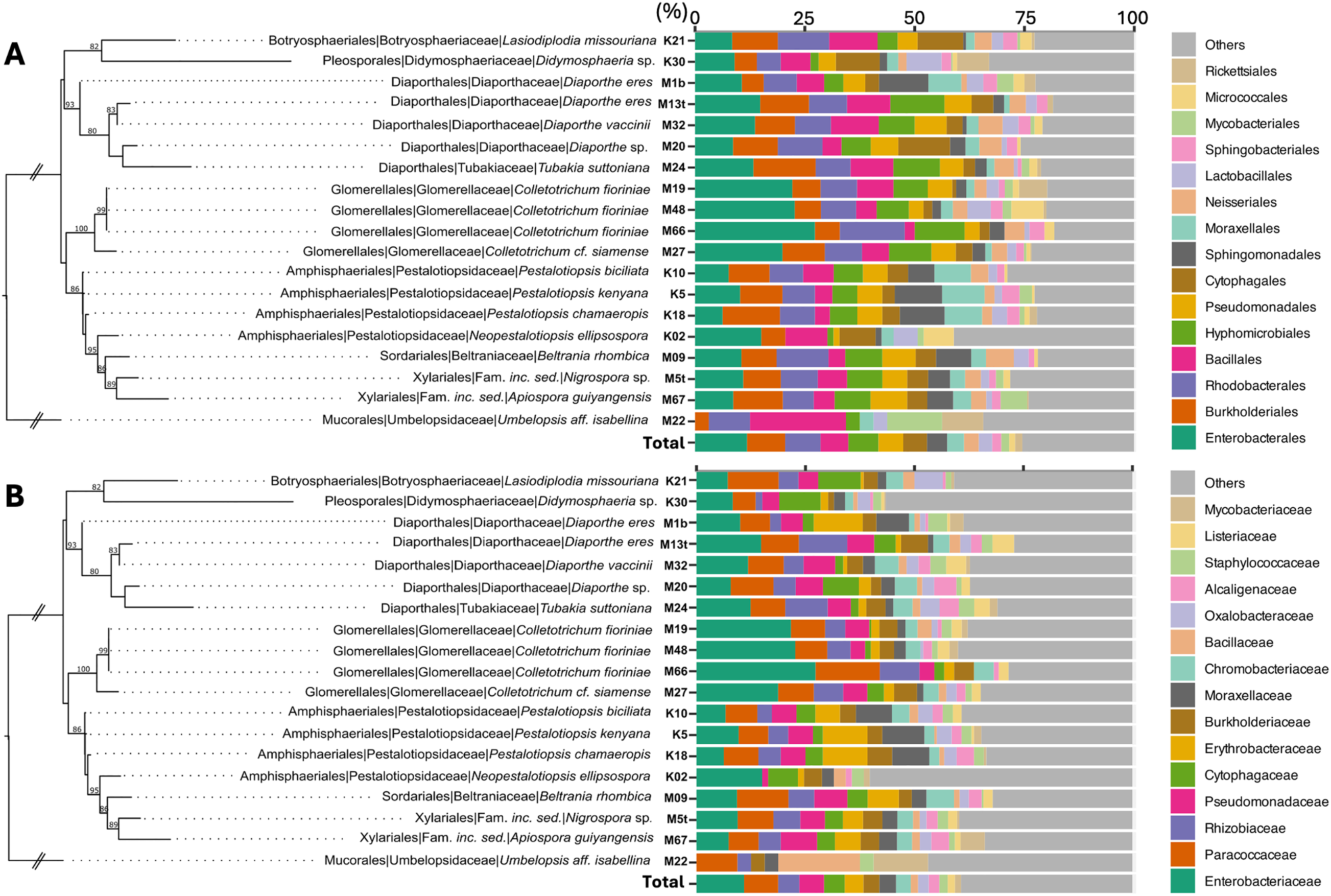
Relative abundance (%) of the top 15 most abundant bacterial orders (A) and families (B) among the fungal isolates. The Maximum Likelihood phylogenetic tree was constructed using concatenated nrDNA ITS, TEF, and TUB sequences. Bootstrap values are shown on nodes. Unidentified taxa were excluded.

Only *Amphisphaeriales*, *Diaporthales*, and *Glomerellales*, had four or more samples each to make inferences about core or resident taxa at the fungal order level (Figure 2, Supplementary Tables S3–S5). The *Amphisphaeriales* had 13 orders, 13 families, and 4 genera classified as core taxa. Examples of core taxa that also had high relative abundances include *Bacillales, Burkholderiales, Hyphomicrobiales*, and *Lactobacillales; Acetobacteraceae, Bacillaceae, Burkholderiaceae*, and *Enterobacteriaceae*; and *Achromobacter, Acinetobacter, Erythrobacter/Porphyrobacter*, and *Klebsiella/Raoultella* (Figures 2 and 3, Supplementary Tables S2–S5). The fungal order *Diaporthales* had 17 orders, 21 families, and 14 genera classified as core (Figure 2, Supplementary Tables S3–S5). Examples of those taxa are *Bacillales, Burkholderiales, Cytophagales*, and *Hyphomicrobiales; Acetobacteraceae, Bacillaceae, Enterobacteraceae*, and *Rhizobiaceae*; and *Coraliihabitans, Erythrobacter/Porphyrobacter, Pseudomonas,* and *Rhizobium/Agrobacterium*. Lastly, *Glomerellales* samples had 15 orders, 17 families, and 12 genera classified as core (Figure 2, Supplementary Tables S3–S5). Examples of core taxa with high relative abundances include *Burkholderiales, Enterobacterales, Hyphomicrobiales*, and *Rhodobacterales; Enterobacteriaceae, Pseudomonadaceae, Paracoccaceae*, and *Rhizobiaceae*; and *Coraliihabitans, Klebsiella/Raoultella, Pseudomonas* and *Rhizobium/Agrobacterium* (Figures 2 and 3, Supplementary Tables S2–S5).

The multilevel pattern analyses made for the best-represented fungal groups indicated significance for some bacterial taxa as potential indicator species. For example, the bacterial family *Streptosporangiaceae* is associated with *Amphisphaeriales* but not *Diaporthales* (*P* = 0.0289) or *Glomerellales* (*P* = 0.0287). For *Glomerellales*, two bacterial taxa, i.e., *Rhizobiaceae* (*P* = 0.0287; *P* = 0.0299) and *Enterobacteriaceae* (*P* = 0.0287; *P* = 0.0299), resulted with significance when compared to *Amphisphaeriales* and *Diaporthales*. Furthermore, the multinomial species test suggested bacterial taxa as potential specialists for *Amphisphaeriales* and *Glomerellales*, i.e., *Moraxellales* and *Sphingomonadales*, and *Enterobacterales* and *Rickettsiales*, respectively (Supplementary Figure S6). The multinomial species test resulted in one specialist bacterial taxon for *Diaporthales* (i.e., *Cytophagales*) and three specialist taxa for *Glomerellales* (i.e., *Enterobacterales, Hyphomicrobiales*, and *Microccocales*) (Supplementary Figure S7). The multinomial species test did not result in any specialist taxa when comparing *Amphisphaeriales* and *Diaporthales*; however, rare and generalist bacteria were represented by *Burkholderiales*, *Enterobacteriales*, and *Pseudomonadales*, among others (Supplementary Figure S8). Finally, the Mantel test confirmed a significant phylogenetic signal in the endohyphal bacterial community (*P* = 0.0026), and the model was deemed robust (0.955) after 10,000 simulations. However, the Procrustes indicated a marginal significance (*P* = 0.042) with a correlation of 0.4689 (Supplementary Figure S9A).

### Viral metagenomic data analyses

#### Alpha diversity

After assembly and taxonomic classification, metagenomic data yielded only 978 contigs that matched viruses (Supplementary Table S6). Sample comparison revealed significant differences in both the number of taxa and contigs. Samples K21 and M67 exhibited similar values, with around 70 taxa and 100 contigs each, while other samples had fewer than 100 contigs and approximately 50 taxa (Supplementary Figure S10A). When classified by fungal host order, *Pleosporales*, *Sordariales*, and *Xylariales* had similar contig counts (∼200), though *Xylariales* displayed the highest taxonomic diversity with around 80 taxa. The taxonomic composition of *Botryosphaeriales* was comparable to these groups, while other orders showed fewer than 50 taxa and 100 contigs. Notably, *Glomerellales*, *Diaporthales*, and *Sordariales* appeared to approach an asymptote (Supplementary Figure S10B). At the family level, samples M5t and M67 accounted for about 200 contigs and nearly 100 taxa. *Botryosphaeriaceae* followed, with around 100 contigs and a slightly lower taxonomic count. Other families contained fewer than 50 taxa. The asymptote was reached by *Diaporthaceae, Glomerellaceae*, and *Pestalotiopsidaceae* (Supplementary Figure S10C). TEM images of the three fungal isolates did not identify clear virus-like particles.

#### Viral community structure, composition, and phylogenetic signal across fungal hosts (metagenomics)

The analysis of the complete dataset determined no significant differences in viral communities at either the order (*F_7,11_* =1.5, *P* = 0.0829) or family (*F_8,10_* = 2.4, *P* = 0.0935) levels, and the communities were homogeneous (*F_7,11_* = 1.5, *P* = 0.1038; *F_8,10_* = 2.4, *P* = 0.1552, respectively). The NMDS stress value suggested a good fit for the model (0.0956), although most of the samples clustered in the lower left panel (Supplementary Figures S11A and S11B). When only the most taxonomically represented fungal samples were retained, the results were similar with no significant differences among fungal orders (*F_2,11_* = 1.55, *P* = 0.0967) or families (*F_2,11_* = 1.55, *P* = 0.1026), or in community homogeneity (*F_2,9_* = 1.92, *P* = 0.2104 for order; and *F_2,9_* = 1.92, *P* = 0.199 for family). The NMDS for the balanced dataset also indicated a good representation of the model (stress value 0.0607). Supplementary Figures S11C and S11D show that sample distances across most orders and families form clear clusters, except for certain samples from *Glomerellales* and *Diaporthales*. *Amphisphaeriales* was the most consistent group.

The viral community composition using metagenomic data revealed that 2 realms (*Duplodnaviria* and *Varidnaviria*) and 2 kingdoms (*Bamfordvirae* and *Heunggongvirae*) were consistently present in 80–100% of the fungal samples (i.e., core); and 2 realms (*Riboviria* and unclassified dsDNA viruses) and 2 kingdoms (unclassified dsDNA viruses and *Pararnavirae*) were present in 50–79% of the samples (i.e., resident) (Figure 4, Supplementary Tables S7 and S8). Interestingly, only about 5% of the detected viral sequences at the realm and kingdom levels were unidentified/unclassified (Figure 5, Supplementary Tables S6–S8). The realms *Duplodnaviria* and *Varidnaviria*, and kingdoms *Bamfordvirae* and *Heunggongvirae* were also the most abundant, with relative abundances ranging from ∼20–60% (Figure 5, Supplementary Tables S6).

**Figure 4.**
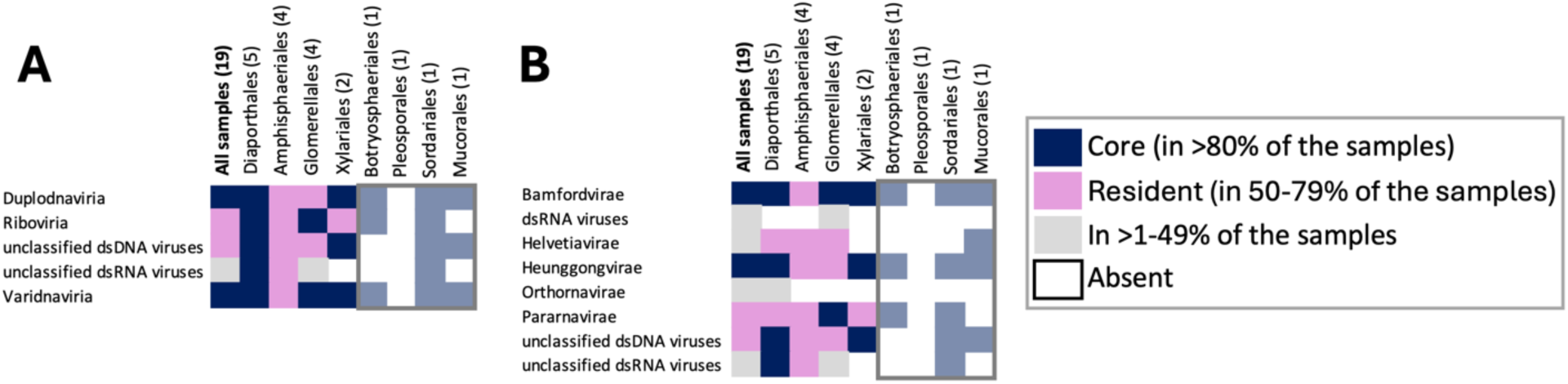
Core and resident viral realms (A) and kingdoms (B) by fungal order, for the metagenomic dataset.

**Figure 5.**
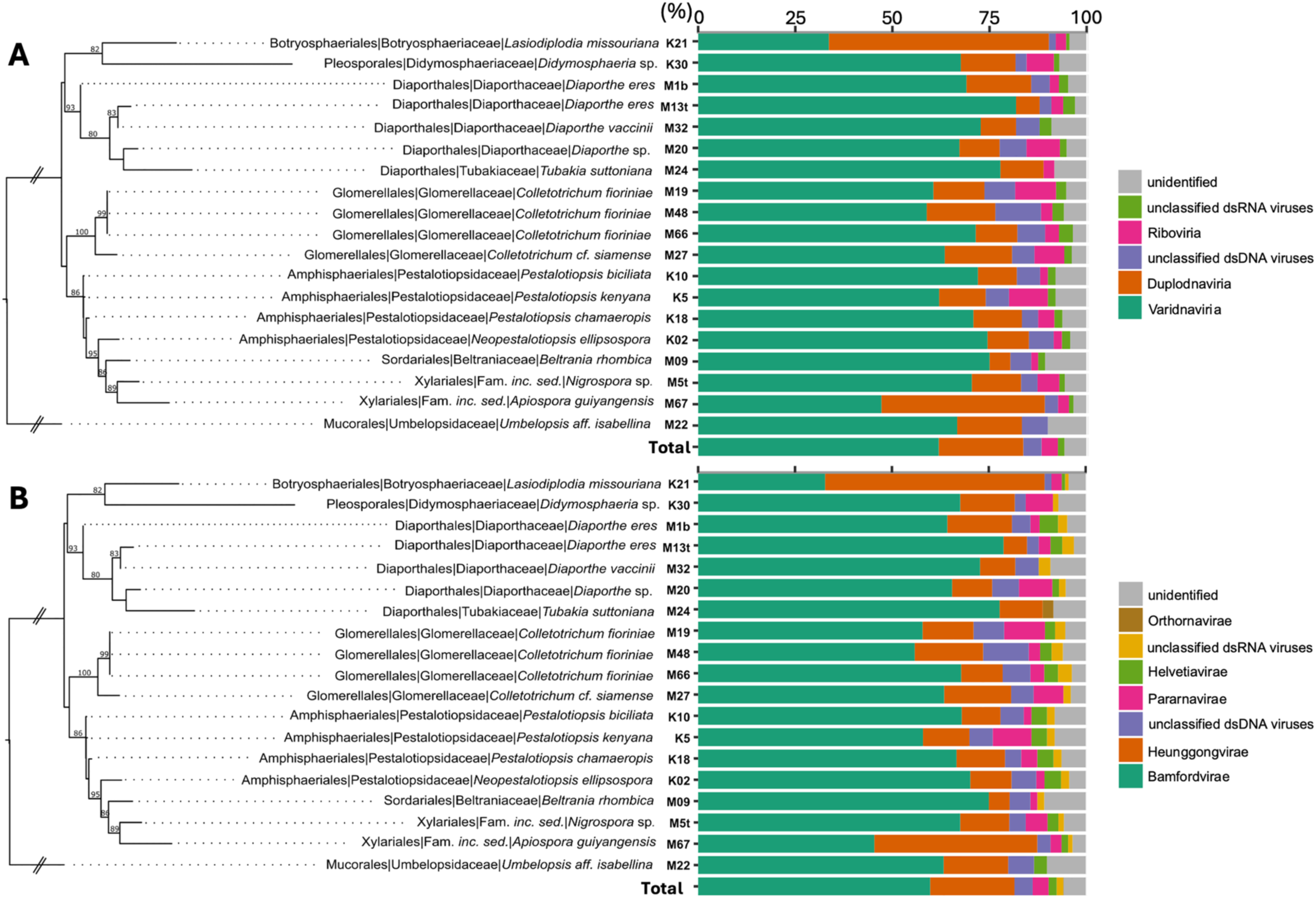
Relative abundance (%) of the viral realms (A) and kingdoms (B) among the fungal isolates, using metagenomic data. The Maximum Likelihood phylogenetic tree was constructed using concatenated nrDNA ITS, TEF, and TUB sequences. Bootstrap values are shown on nodes.

Only *Amphisphaeriales*, *Diaporthales*, and *Glomerellales*, had four or more samples each to make inferences about core or resident taxa (Figure 4, Supplementary Tables S7 and S8). The *Amphisphaeriales* had no taxa classified as core, but 5 realms and 6 kingdoms classified as resident. The resident realms, from the most abundant to the least, were *Varidnaviria*, *Duplodnaviria*, unclassified dsDNA viruses, *Riboviria*, and unclassified dsRNA viruses; and kingdoms *Bamfordvirae*, *Heunggongvirae*, unclassified dsDNA viruses, *Pararnavirae, Helvetiavirae*, and unclassified dsRNA viruses (Figures 4 and 5, Supplementary Tables S6–S8). The fungal order *Diaporthales* had 5 realms and 4 kingdoms classified as core, and no resident realms but 2 resident. The core realms, from the most abundant to the least, were *Varidnaviria*, *Duplodnaviria*, unclassified dsDNA viruses, *Riboviria*, and unclassified dsRNA viruses; and kingdoms *Bamfordvirae*, *Heunggongvirae*, unclassified dsDNA viruses, and unclassified dsRNA viruses (Figures 4 and 5, Supplementary Tables S6–S8). The resident kingdoms were *Helvetiavirae* and *Pararnavirae*. Lastly, *Glomerellales* had 2 core realms (*Riboviria* and *Varidnaviria*), 2 core kingdoms (*Bamfordvirae* and Pararnavirae), 2 resident realms (*Duplodnaviria* and unclassified dsDNA viruses) and 3 resident kingdoms (*Helvetiavirae*, *Heunggongvirae*, and unclassified dsDNA viruses) (Figure 4, Supplementary Tables S7 and S8).

The multilevel pattern analysis only detected two taxa, i.e., *Duplodnaviria* (*Uroviricota, Caudoviricetes*) and *Varidnaviria* (*Nucleocytoviricota, Megaviricetes*), resulting in a significant association (*P* = 0.0268 for each) with its fungal host *Amphisphaeriales* when compared to *Diaporthales*. Similarly, the multinomial test found generalist and rare viral taxa for *Amphisphaeriales*, *Diaporthales*, and *Glomerellales* when compared to each other; however, no particular or specialist taxa were detected for any of those orders (Supplementary Figure S12– S14). Viral taxa for all the analyses, classified as generalist belong to *Varidnaviria* and *Duplodnaviria*, while unclassified dsRNA or *Riboviria* were classified as rare. An evaluation of the phylogenetic signal for the full dataset yielded a negative result (0.3729). However, when the analysis was restricted to the three best-represented fungal orders, the test suggested a significant phylogenetic signal (*P* = 0.0016) with a robust model (0.95) after 10,000 simulations. The phylogenetic signal was supported by the Procrustes analysis, which found a correlation between the viral communities and their fungal hosts (correlation 0.9431, *P* = 0.001) (Supplementary Figure S9B).

### Viral metatranscriptomic data analyses

#### Alpha diversity

After assembly and taxonomic classification, the metatranscriptomic data recorded a total of 8,245 transcripts that matched viruses (Supplementary Table S9). Most samples contributed over 500 transcripts, except for M9, M32, and M48, which had around 200. However, extrapolated data predicted up to 1,000 transcripts and nearly 400 taxa. On average, the number of taxa across samples was around 200 (Supplementary Figure S15A). At the fungal order level, *Sordariales* exhibited the highest diversity, with 300 taxa across 1,000–2,500 transcripts. *Amphisphaeriales* accounted for fewer than 100 taxa and 1,000 transcripts, while other orders presented around 200 taxa and close to 1,000 transcripts. Extrapolated data suggested that most groups could reach approximately 400 taxa (Supplementary Figure S15B). At the family level, *Diaporthaceae, Glomerellaceae*, and *Pestalotiopsidaceae* showed the highest number of transcripts (∼1,500) and taxa (>200). Most other families reached around 500 transcripts and 200 taxa, while *Beltraniaceae* had the fewest (<1,000 transcripts and <100 taxa). Extrapolated data indicated the potential for all groups to have higher reads and taxon counts, with *Diaporthaceae* the only family approaching an asymptote (Supplementary Figure S15C).

#### Viral community structure, composition, and phylogenetic signal across fungal hosts (metatranscriptomics)

Beta diversity analysis revealed differences in viral communities at both the fungal order (*F_7,11_* = 9.48, *P* = 0.0006) and family (*F_9,9_* = 10.8, *P* = 0.0007) levels using the full dataset. Community dispersion showed heterogeneity for both taxonomic levels (order: *F_7,11_* = 13.9, *P* = 0.0002; family *F_9,9_* = 10.82, *P* = 0.0012). The NMDS resulted in a good fit (stress value 0.0902); nonetheless, no clear clusters were observed at either the fungal order or family level (Supplementary Figures S16A and S16B). In contrast, with the balanced dataset, differences were found at the fungal order level (*F_2,11_* = 1.87, *P* = 0.0259), but not at the family level (*F_4,11_* = 1.06, *P* = 0.3807). Community dispersion showed homogeneity at the order level (*F_2,9_* = 3.74, *P* = 0.0557) but heterogeneity at the family level (*F_4,7_* = 9.03, *P* = 0.0041). The NMDS for the balanced dataset indicated a good fit (stress value 0.0621). Supplementary Figure S16C shows that samples from *Glomerellales* (M19 and M48) cluster closely, while *Amphisphaeriales* form a distinct group. *Diaporthales*, on the other hand, appear to form two groups in the upper quadrants. A similar pattern is observed at the family level (Supplementary Figure S16D).

The viral community composition using metatranscriptomic data revealed no realms or kingdoms consistently present in 80–100% of the fungal samples (i.e., core); but 4 realms (*Duplodnaviria*, *Riboviria*, *Varidnaviria*, and unclassified dsDNA viruses) and several kingdoms (*Bamfordvirae*, *Helvetiavirae*, *Heunggongvirae*, *Orthornavirae, Pararnavirae*, and unclassified dsRNA and dsDNA viruses) were present in 50–79% of the samples (i.e., resident) (Figure 6 Supplementary Tables S10 and S11). About 42% of the detected viral sequences at the realm and kingdom levels were unidentified/unclassified (Figure 7, Supplementary Table S9). The realms *Duplodnaviria* and *Varidnaviria*, and kingdoms *Bamfordvirae* and *Heunggongvirae* were also the most abundant, and consistent with the metagenomic data, with relative abundances of ∼25% (Figure 7, Supplementary Table S9).

**Figure 6.**
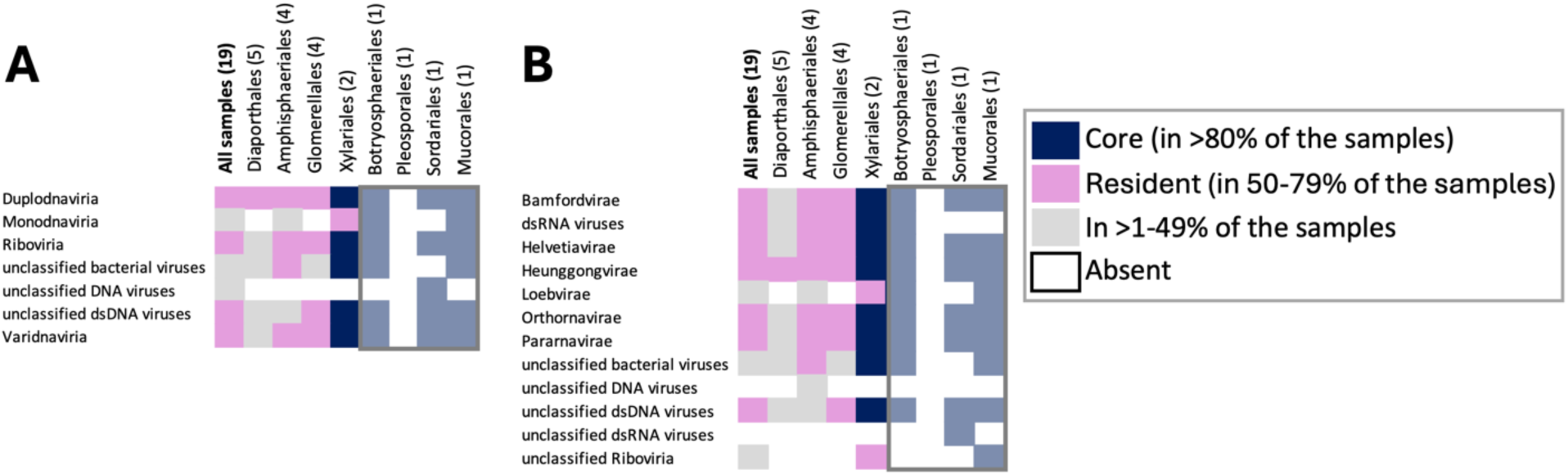
Core and resident viral realms (A) and kingdoms (B) by fungal order, for the metatranscriptomic dataset.

**Figure 7.**
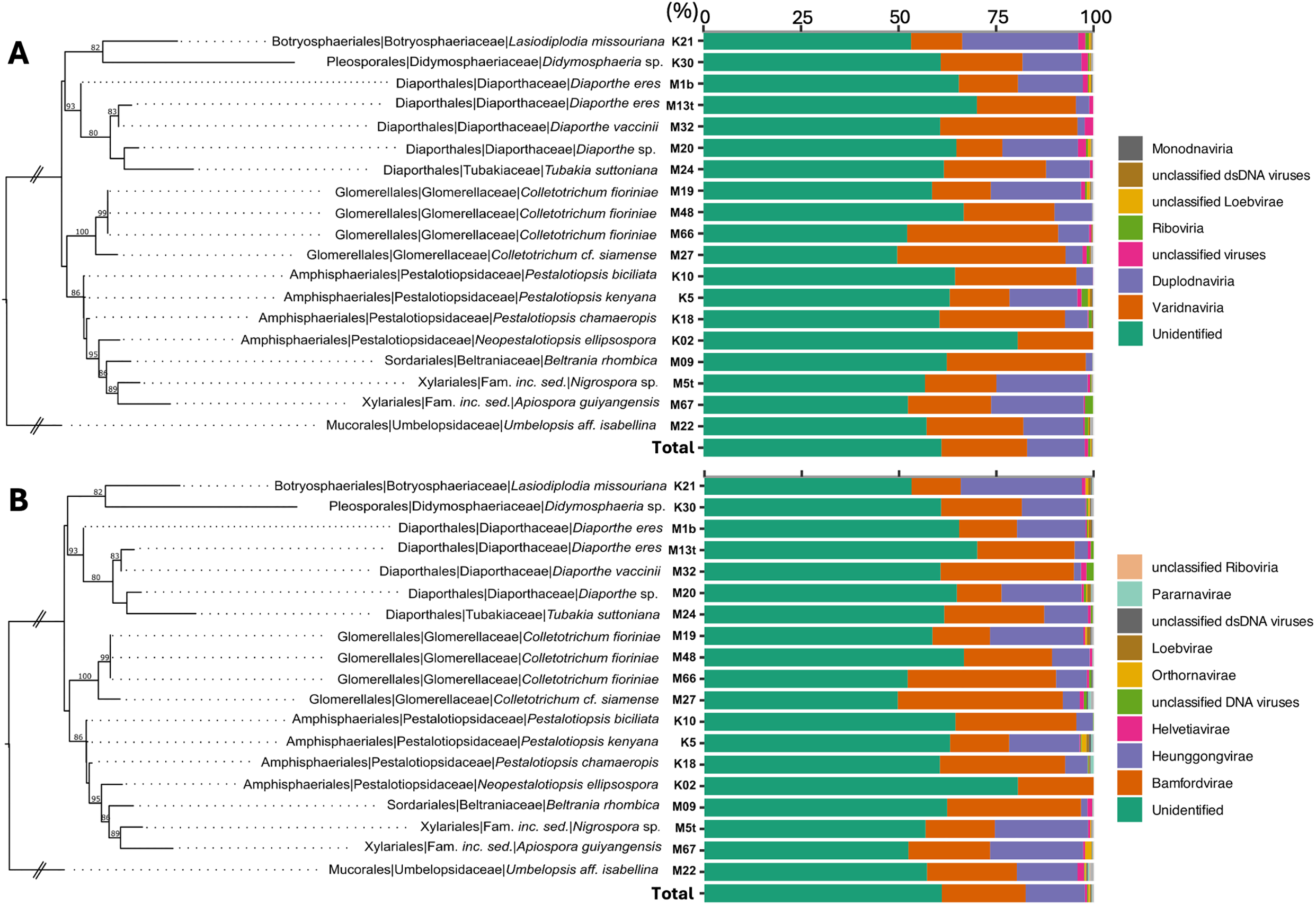
Relative abundance of the viral realms (A) and kingdoms (B) among the fungal isolates, using metatranscriptomic data. The Maximum Likelihood phylogenetic tree was constructed using concatenated nrDNA ITS, TEF, and TUB sequences. Bootstrap values are shown on nodes.

As mentioned before, only *Amphisphaeriales*, *Diaporthales*, and *Glomerellales*, had four or more samples each to make inferences about core or resident taxa (Figure 6, Supplementary Tables S10 and S11). None of the three fungal orders had viral realms or kingdoms classified as core. The *Amphisphaeriales* had 4 realms and 7 kingdoms classified as resident. The most abundant resident realm was *Duplodnaviria* (∼18%), followed by others with abundances <0.5%; and the most abundant kingdoms were *Bamfordvirae* (∼26%) and *Heunggongvirae* (∼18%), followed by *Helvetiavirae, Pararnavirae*, unclassified dsRNA and dsDNA viruses, and others with relative abundances of <1.5% (Figures 6 and 7, Supplementary Tables S9–S11). About 54% of the viral transcripts were unclassified at the realm or kingdom level. The fungal order *Diaporthales* had one realm (*Duplodnaviria*) and one kingdom (*Heunggongvirae*) with relative abundances of ∼10% (Figure 7, Supplementary Table S9). M24 (*Tubakia suttoniana*) and M32 (*Diaporthe vaccinii*) were particularly abundant (∼44–75%) in *Varidanaviria* (Supplementary Table S9). About 73% of the viral transcripts were unclassified at the realm or kingdom level. Lastly, *Glomerellales* also had 4 resident realms and 7 resident kingdoms. The most abundant resident realms were *Varidnaviria* (∼35%) and *Duplodnaviria* (∼23%), followed by others with abundances <2%; and the most abundant kingdoms were *Bamfordvirae* (∼33%) and *Heunggongvirae* (∼23%), followed by others with relative abundances of <2% (Figures 6 and 7, Supplementary Tables S9–S11). About 39% of the viral transcripts were unclassified at the realm or kingdom level.

After performing the multilevel pattern analyses for the best-represented fungal groups, viral taxa related to *Uroviricota* (*Duplodnaviria*) and *Nucleocytoviricota* (*Varidnaviria*) were found to be significant for *Glomerellales* (*P* = 0.029) when compared to *Amphisphaeriales*. Other taxa, *Naldaviricetes* (*Varidnaviria*) resulted significant (*P* = 0.0268) when the *Glomerellales* was compared with *Diaporthales* (Supplementary Figure S17). However, no specific viral taxa were found for *Amphisphaeriales* or *Diaporthales*. In contrast, the multinomial test found specialist taxa for *Diaporthales* when compared to *Amphisphaeriales*; however, the specialist taxa belong to unidentified viral taxa. Nonetheless, when compared to *Glomerellales*, signals were mainly related to *Varidnaviria* (*Megaviricetes*) (Supplementary Figures S18 and S19). The Mantel test to assess phylogenetic signal with the entire metatranscriptomic dataset indicated a negative result (*P* = 0.3999). However, a marginal difference was observed with the balanced dataset (*P* = 0.0539) after 10,000 iterations. Both analyses produced robust models (0.959). Lastly, the Procrustes analysis found significance for the association of the viral metatranscriptomic community and its fungal hosts (*P* = 0.001, correlation 0.8033) (Supplementary Figure S9C).

## DISCUSSION

### Host specialization and community structure

Significant differences in beta diversity among fungal orders and families, combined with the presence of core bacterial and viral taxa, suggest a host-specific endohyphal community associated with the host fungus. This is further corroborated by the strong phylogenetic signal observed, particularly in bacterial communities, indicating that host association is influenced by phylogenetic relatedness. Such relatedness, in turn, plays a crucial role in shaping the composition of these microbial communities [82,83]. Notable bacterial taxa identified as indicator species or specialists for specific fungal groups include *Moraxellales, Sphingomonadales*, and *Streptosporangiaceae* for *Amphisphaeriales*; *Enterobacterales* (e.g., *Enterobacteraceae*), *Hyphomicrobiales* (e.g., *Rhizobiaceae*), and *Micrococcales* for *Glomerellales*; and *Cytophagales* for *Diaporthales*. While many additional taxa were classified as core, indicator, or specialist bacterial taxa were encompassed within this core category.

In contrast, the mixed results for viral communities highlight the complexity of these interactions. Overall, there was no concordance between the core taxa classification, and the multilevel pattern (indicator) and multinomial (specialist) tests. Furthermore, although viral communities did not show significant differences across fungal orders and families in the full dataset, the significant phylogenetic signal in the balanced dataset suggests an underlying pattern of association when focusing on specific fungal orders. For example, some core, indicator, or specialist viral taxa may be closely associated with endohyphal bacteria or integrated within fungal genomes. Notably, *Caudoviricetes* (*Duplodnaviria*, *Uroviricota*, *Heunggongvirae*), which includes many bacteriophages, and *Megaviricetes* (*Varidnaviria*, *Nucleocytoviricota*, *Bamfordvirae*), found as possible endogenous elements in fungal genomes, illustrate these associations [49,84]. There are also other examples of *Nucleocytoviricota* in protists, algae, and arthropods, among other eukaryotes [85,86]. Therefore, the lack of significant differences in the full dataset; mixed results in core, indicator, and specialist taxa; but significant phylogenetic signal suggest that other factors, such as the environment, mode of virus transmission, or molecular host-virus interactions [10,26–28,45,46,87–89], might also influence viral community composition and host specialization.

### Endohyphal bacterial communities

In the present work, a great diversity and abundance of endohyphal bacteria were found among the fungi sampled, supporting previous studies [11,24,90,91]. Additionally, the lack of an asymptote in the taxa accumulation curves suggests that much greater diversity remains to be characterized. The detection of core bacterial taxa in the fungal samples, suggests a level of specialization that could be vital for the functioning and stability of these symbiotic relationships [92–96]. Some noteworthy bacteria that were found belong to the *Bacillales, Burkholderiales, Enterobacterales, Hyphomicrobiales*, and *Pseudomonadales*, which are known to play a role in fungal phenotype and symbiosis establishment. For example, some endohyphal *Enterobacter* species can increase the expression of important toxins in fungi (e.g., fumonisin in *Fusarium fujikuroi*) [20] or change their plant pathogenicity by altering genes in their hosts [97]. Members of the *Burkholderiaceae*, which have been reported as abundant in *Mucoromycota* [98], play a significant role in modulating rhizoxin production [99]. This compound, previously attributed to the pathogenic fungus *Rhizopus*, is actually produced by endohyphal *Burkholderia* spp.

*Burkholderiaceae*, in general, has consistently been found in many groups of fungi [95]. *Rhizobium radiobacter* (*Hyphomicrobiales*) is related to the successful endophytic colonization of the basidiomycete *Piriformospora indica* as well as to the improvement of the systemic protection of different crops against bacterial pathogens such as *Xantomonas traslucens* or *Pseudomonas syringae* [23,24,100]. Spores from *Glomeromycota* (arbuscular mycorrhizal fungi) associated with *Bacillus*, *Pseudomonas,* and *Rhizobium* bacteria suggest a recruited microbiota to increase eventual fungal dispersal and colonization success [101]. In our study, some bacterial orders detected through the metagenomic data have not been reported before as endohyphal (e.g., *Lactobacillales, Micrococcales, Moraxellales, Rhodospirillales*, or *Vibrionales*) and therefore, future studies could focus on elucidating the functional contributions of these newly identified bacterial taxa to fungal metabolism and fitness.

### Endohyphal viral communities

In contrast to the diversity of endohyphal bacteria, the viral sequence diversity in our study was lower. This is supported by taxa accumulation curves, which reached an asymptote for some fungal hosts. For instance, we only identified *Bamfordvirae* (*Varidnaviria*) and *Heunggongvirae* (*Duplodnaviria*), both with dsDNA genomes, as core taxa across all fungal orders in the metagenomic data. No core taxa were identified in the metatranscriptomic data. These findings contradict other studies that reported *Riboviria* (i.e., dsRNA viruses) and *Monodnaviria* as the most abundant mycoviral realms [7,10]. In our study, *Riboviria* sequences were detected in approximately 4% and 1% of the metagenomic and metatranscriptomic datasets, respectively, while *Monodnaviria* was absent in the metagenomic dataset and present at a negligible level (∼0.04%) in the metatranscriptomic dataset. In addition, we expected to find more dsRNA mycoviruses, but we only detected a few contigs or transcripts of dsRNA mycoviruses (e.g., *Diplodia scorbiculata* RNA virus 1–like and other unidentified dsRNA viruses). Despite these differences, roughly 40% of the viral transcripts remained unclassified, highlighting the challenges in virome studies and the need for expanded and improved viral taxonomic databases [102–104].

The *Caudoviricetes* (*Duplodnaviria*, *Uroviricota*, *Heunggongvirae*), which are abundant across all fungal orders in our study and include bacteriophages, have the potential to influence fungal phenotypes and genetic architecture. Phages can indirectly affect fungi through interactions with bacterial intermediaries and horizontal gene transfer, playing a critical role in shaping fungal phenotypes and microbial ecosystems. For example, phages can transfer DNA between bacteria and, in some cases, to fungi, a process known as bacteriophage-mediated gene transfer, which, in consequence, can have significant ecological and evolutionary implications [105].

Bacteriophage-mediated gene transfer has been shown to influence fungal colonization of plant roots [105], fungal degradation processes [95], and the production of fungal secondary metabolites, such as terpenoids, which serve diverse ecological functions [106]. These metabolites can kill nematodes and arthropods, attract animals to facilitate spore dissemination, and mediate communication between fungi and bacteria [106]. Additionally, phages can influence fungi indirectly by affecting bacterial populations through bacterial lysis and community dynamics. Phage-mediated bacterial lysis releases key nutrients—such as nitrogen, phosphorus, and carbon—into the environment, promoting fungal growth and metabolism [107,108]. In the rhizosphere, phages infecting nitrogen-fixing bacteria, such as *Rhizobium*, can disrupt nutrient cycling [109–111], thereby reducing resources available to fungi. Furthermore, phages can modulate the production of bacterial secondary metabolites, which often play a key role in bacterial-fungal interactions. For instance, phages infecting *Bacillus* species may enhance the production of antifungal compounds like iturin A, thereby altering fungal pathogen suppression [112,113].

*Mimiviridae* and *Phycodnaviridae* (*Varidnaviria*, *Bamfordvirae*, *Megaviricetes*) were the most abundant viral families. These families have been reported mostly in amoebae, algae, and bacteria. Although direct infections of fungi by members of *Nucleocytoviricota* (giant viruses) are not well-documented [49,84,114], there is evidence of interactions through endogenous viral elements and environmental viromes. For example, a significant 1.5-Mb endogenous viral region, related to the family *Asfarviridae* within *Nucleocytoviricota*, was discovered in the genome of the arbuscular mycorrhizal fungus *Rhizophagus irregularis* [49]. This suggests ancient viral integration events that may influence fungal evolution and genome architecture.

Studies have also identified viral sequences related to *Nucleocytoviricota* in various environments, including freshwater ecosystems [115]. These findings indicate the presence of giant viruses in habitats where fungi are prevalent, suggesting potential interactions.

Additionally, research has uncovered complex genomes of early nucleocytoviruses through ancient endogenous viral elements in diverse eukaryotic lineages, including fungi [116]. This highlights the role of giant viruses in horizontal gene transfer and their potential impact on the evolution of fungal genomes.

### Limitations and future directions

The lack of visible bacterial-like cells (in TEM) within isolate M67 could indicate the sectioned hyphae had not yet been invaded by the bacteria or had lost the bacteria, possibly via repeated subculturing [13]. However, the presence of visible bacteria within the hyphae of isolates M5t and K21 provides direct evidence in support of the sequencing analysis. In contrast, within the isolates investigated, no clear virus-like particles were observed, suggesting that mycoviruses occur at low titers, at least within the cultures studied, which supports the relatively low number of contigs and transcripts found in the metagenomic and metatranscriptomic assemblies.

Additional studies that concentrate virus particles from larger cultures or specific PCR amplifications using primers designed from the identified scaffolds are needed to further investigate this topic.

Additional limitations should be acknowledged. One is the high percentage of unclassified taxa, which may affect interpretative power, underscoring the need for expanded reference databases, especially for viruses [102–104]. Notably, many of the specialist taxa (resulting from the multinomial classification) are unidentified. Furthermore, our sample size for certain fungal orders was limited, which may have impacted the robustness of phylogenetic and community analyses. Expanding the sample size for underrepresented fungal orders and including additional orders will strengthen inferences about the phylogenetic signal and host specialization, especially for endohyphal bacteria, which did not reach an asymptote in the species accumulation curve in our metagenomic data. We also did not consistently assess the presence of viral or bacterial sequences inserted into fungal genome assemblies (e.g., endogenous elements) or determine whether these insertions were recent or ancestral events [47–49]. Investigating viral and bacterial sequences integrated into fungal or eukaryotic genomes can provide compelling insights into evolutionary and functional impacts [47,117,118]. Lastly, the reliance on high-throughput sequencing may potentially introduce biases, such as uneven detection of RNA versus DNA viruses [119].

## CONCLUSIONS

Our study is one of the first that have used phylogenetic signals to uncover potential host specialization in endohyphal microbial communities and provide a framework for further exploring co-evolutionary dynamics, enriching our knowledge of multi-partite host-microbe interactions and their evolutionary implications. In this study, we showed phylogenetic and ecological relationships between three fungal orders in the *Ascomycota* (*Sordariomycetes*) and their bacterial and viral inhabitants, revealing intriguing patterns of potential host specialization and microbial interactions. The significant phylogenetic signal and the identification of core bacterial taxa across fungal samples and orders provide evidence for specialized fungal-microbe associations, underscoring the critical roles these communities play in fungal and holobiont biology and ecosystem functioning. In contrast, viral community analyses presented mixed results, with significant phylogenetic signals only in specific datasets, highlighting the complexity of fungal-viral interactions. The detection of *Caudoviricetes* (bacteriophages) and *Megaviricetes* (giant viruses), suggests that horizontal gene transfer and genomic integration may be impacting the phenotype and genomes of the fungi sampled. However, key limitations, including a high percentage of unclassified taxa and limited sample size for some fungal groups, emphasize the need for expanded reference databases, broader sampling efforts, and deeper functional investigations. Future research should focus on elucidating the functional contributions of newly identified or unidentified bacterial and viral taxa, exploring endogenous viral and bacterial elements in fungal genomes, and examining the ecological and functional roles of these microbial communities in the endophytic niche and diverse environments. This study provides a foundation for understanding fungal-microbe interactions, with implications for microbial ecology, evolutionary biology, and applied microbiome research.

## LIST OF ABBREVIATIONS

ANOVA: analysis of variance
BLAST: Basic Local Alignment Search Tool
BSA: bovine serum albumin
CeNAT-CONARE: Centro Nacional de Alta Tecnología, Consejo Nacional de Rectores
dbRDA: distance-based redundancy analysis
DMSO: dimethyl sulfoxide
dsRNA/DNA: double-stranded RNA/DNA
HPC: high-performance computing
ITS nrDNA: internal transcribed spacer regions of the nuclear ribosomal DNA
MEB: malt extract broth
NCBI RefSeq: National Center for Biotechnology Information, Reference Sequence Database
NMDS: non-metric multidimensional scaling
PCR: polymerase chain reaction
PDA: potato dextrose agar
PERMANOVA: permutational multivariate analysis of variance
TEF: translation-elongation factor 1-α
TEM: transmission electron microscopy
TUB: β-tubulin

## DECLARATIONS

### Availability of data and material

Supplementary tables and figures supporting the results and conclusions of this article are available in the Zenodo repository (https://doi.org/10.5281/zenodo.14847255). Raw .fastq files for metagenomic and metatranscriptomic data are deposited in GenBank’s Sequence Read Archive (SRA) under BioProject PRJNA1221291. GenBank accession numbers for newly generated sequences (ITS nrDNA, TEF, and TUB) are indicated in Supplementary Table S1.

### Authors’ contributions

Conception: PC, MC, JNC; design of the work: PC, JNC, ESL; data acquisition: PC, MB, NG, AD, JK; data analysis: PC, ESL, MA, NG, MB, JK, JNC; data interpretation: PC, ESL, MA, JK, JNC, MA, MC; drafted the written work: ESL, PC; revised the written work: PC, ESL, MC, JNC, JK, MB, NG, AD, MA.

## Acknowledgments

We would like to acknowledge Bowie State University students from the Course-based Undergraduate Research Experience (CURE) course Microbiology BIOL309 (Spring 2024) for their assistance in DNA and RNA extraction from four fungal samples.

